# Longitudinal survey reveals delayed effects of forager gene expression on stingless bee colony health

**DOI:** 10.1101/2020.10.13.337527

**Authors:** Lílian Caesar, Anelise Martins Correa Lopes, Jefferson Nunes Radaeski, Soraia Girardi Bauermann, Enéas Ricardo Konzen, Jean-François Pombert, Aroni Sattler, Betina Blochtein, Airton Torres Carvalho, Karen Luisa Haag

## Abstract

Bee populations are declining globally due to different environmental stressors, such as pathogens, malnutrition, and agrochemicals. Brazil is the home of hundreds of stingless bee species, some of them now considered endangered, though very little is known about the impact of disease on native bees. In Southern Brazil the endangered stingless bee *Melipona quadrifasciata* is affected by an annual syndrome that causes sudden death of workers, eventually leading colonies to collapse. Although novel viruses were found in foragers from diseased colonies, none has been consistently implicated in the outbreak. Here we used transcriptomics in combination with an integrative longitudinal survey on managed colonies to identify predictors for preventing *M. quadrifasciata* colony failures. We found that key genes related to xenobiotic metabolization, nutrition and immune responses are downregulated in foragers from colonies that became diseased three months later. The period that preceded the outbreak was marked by pronounced forager weight loss as well as behavioural changes. Our findings support the proposition that worldwide bee mortality is influenced by a combination of diverse sublethal factors, and increase awareness of the long-term effects of genetic diversity erosion in stingless bee species, which enhances their vulnerability to environmental stressors.

## Introduction

The worldwide decrease in bee populations observed in the last decade is a matter of great concern. Some colony losses are explained by the presence of pathogens and other infectious agents [1,2], but several additional interacting stressors, such as habitat loss, malnutrition, agrochemicals and colony management practices are known to reduce the fitness of bee populations [3]. Theoretical studies indicate that although multifactorial stresses may cause colony failure, it is most likely a result of a critical level of stress from the accumulation of sublethal factors [4].

One of the intrinsic properties of sublethal factors is their delayed effect on organismal fitness. In eusocial bees, where individual fitness is achieved indirectly by cooperation, sublethal factors characteristically compromise the ability of non-reproductive females to perform their regular tasks [5]. For example, honey bee workers reared in pollen-stressed colonies show normal development, but as adults are less likely to waggle dance, and precisely inform food location [6]. Exposure of larvae to sublethal doses of pesticides induces multiple changes in gene expression, leading to developmental and adult behavioural changes that weaken bee colonies [5,7]. Besides interacting with each other, environmental stressors are modulated by endogenous factors that are ultimately linked to the bee’s genetic background [8]. Some bee management practices, including colony translocation, may lead to genetic homogenization, loss of genetic diversity, and the breakdown of local adaptations, finally impacting their capacity to respond to stressors [9,10].

The stingless bee *M. quadrifasciata* is one of the most extensively managed species and the object of intensive trading in Southern Brazil, where its nests virtually disappeared from nature [11]. In this region, the practice of colony division has been performed for decades and became more common with trade intensification [12], which is now threatened by a syndrome that annually occurs in late summer, often leading to colony collapse [13,14]. Some worker bees from affected colonies show neurologic symptoms, such as tremors and paralysis, suggesting the implication of viruses in the syndrome. However, in spite of having found novel viruses associated with symptomatic bees, such as dicistroviruses, which cause similar symptoms in *A. mellifera* [15], no virus [14] or other pathogen [13] was found consistently associated with bees from diseased colonies. Here we report an integrative and longitudinal study designed to uncover the causes underlying the annual syndrome of *M. quadrifasciata*. In addition to exploratory transcriptomic analyses, a temporal survey was conducted on three pairs of mother-daughter colonies kept in two separate localities, in order to evaluate the contribution of both genetic and environmental factors to the syndrome manifestation, by measuring individual- and colony-level traits.

## Materials and Methods

### (a) Transcriptome analyses

Three *M. quadrifasciata* colonies were sampled for exploratory transcriptomic analyses aiming to identify differentially expressed genes (DEGs) during the syndrome, and to select candidate genes for further analyses. Two colonies (D1 and D2) presented symptoms during the yearly outbreak, while the third (H) remained healthy and was sampled after the outbreak period. RNA of three pooled bees from each colony was purified by polyA-tail selection, followed by library construction using TruSeq Stranded mRNA Library Prep Kit (Illumina, USA). Single-end sequencing (read length = 150 nt) was performed on an Illumina NextSeq instrument. Low quality reads were removed with Trimmomatic v.0.36 [16], and mapped onto *M. quadrifasciata* genome (GenBank: GCA_001276565.1) with GSNAP v. 2018-07-04 [17]. Gene expression was estimated with the *depth* command from Samtools v.1.3.1 [18], and DEGs were recovered with custom Perl scripts, after a normalization step (Available at https://github.com/liliancaesar/Publication_scripts/tree/main/2020_Longitudinal_survey/Transcriptome_analyses). Functional enrichment analysis was performed with g:Profiler [19]. Detailed information on the samples, nucleic acid extraction and transcriptome analyses are provided as electronic supplementary material (text S1). Three DEGs known for their roles in bee health were selected for RT-qPCR assays.

### (b) Monitoring bee colonies and forager relative gene expression

To monitor changes of key biological features, observations were made in *M. quadrifasciata* mother-daughter (MD) colonies under semi-controlled conditions during six months in two localities. Three colonies (named BP1, BP2 and BP3) were kept in a small agricultural property where agrochemicals are regularly used. Daughter colonies obtained by division from each BP colony (named PA1, PA2 and PA3), were translocated to the vicinity of a secondary forest, located about 70 km away from BP. Colonies were equipped with a datalogger device (ONSET, Brazil) to record within-hive temperature and humidity every six hours. Downstream analyses were conducted with the daily lowest temperature (t) and the highest humidity (h), or with a variable called delta (Δ = maximum value - minimum value). Aliquots of pollen stored inside colonies were collected every month from the experimental hives or from nearby colonies in each meliponary. Around 500 pollen grains were identified at the family, genus, or species level, using reference material from pollen libraries and the literature (see electronic supplementary material, text S1). Every month five foragers (figure 1A) were collected, weighted, and stored at −80°C until RNA extraction for RT-qPCR.

**Figure 1:**
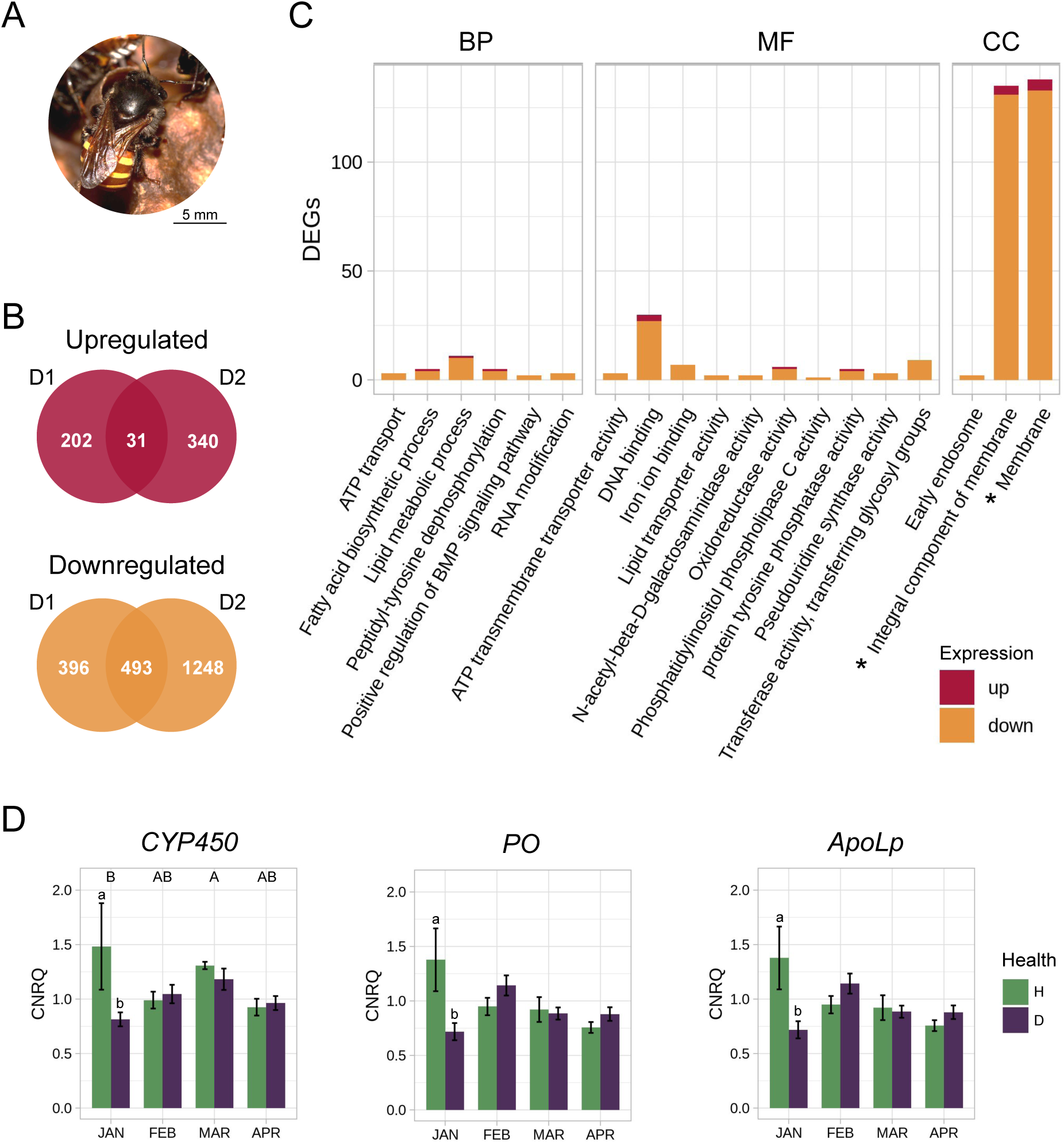
(A) *Melipona quadrifasciata* worker bee (Fototeca Cristiano Menezes, http://www.splink.org.br/search?lang=en&collectioncode=FCM). (B) Venn diagram showing differentially expressed genes (DEGs) up or downregulated in foragers from diseased colonies (D1 or D2). (C) Functional overview of DEGs based on gene ontology annotation (*P <* 1), with the asterisk indicating GO terms enriched in the DEGs (*P* < 0.05). Bars are sorted according to biological processes (BP), molecular function (MF) and cellular component (CC). (D) Barplot comparing relative expression (CNRQ) of *CYP450*, *PO* and *ApoLp* in foragers from colonies that remained healthy (H), with those that showed signs of disease (D) during the outbreak period. Bars indicate the standard error; uppercase letters indicate significant differences among months, and lowercase letters indicate significant differences between healthy and unhealthy colonies within months, according to Tukey’s test (*P* < 0.05).

Three DEGs were selected for monitoring forager bee expression patterns during the survey, namely putative mitochondrial *cytochrome P450* (*CYP450;* WN51_04136), *phenoloxidase* (*PO;* WN51_02761) and *apolipophorin* (*ApoLp*; WN51_14077). Primers based on each respective gene sequence were designed with Primer3 from Geneious R11 [20] (electronic supplementary material, table S1). Actin (*act*) and 40S ribosomal protein S5 (*rps5*) were used as references for gene expression normalization [21–23]. StepOnePlus Real-Time PCR System (Applied Biosystems, USA) was used for the RT-qPCR assays, and primer amplification efficiency was calculated with qBASE+ software [24] from the slope of a five-point 1:10 serial dilution of calibrator cDNA samples. Experimental setup of qPCR involved sample maximization method, three technical replicates for each sample, and inter-run calibrator samples [24]. The default of qBASE+ pipeline was used to calculate relative gene expression, and presented in the form of calibrated normalized relative quantities (CNRQ) [24]. Detailed information on sample processing and RT-qPCR quantifications is provided as electronic supplementary data (text S1).

### (c) Statistical analyses

All analyses were conducted in R version 3.6.3 [25] and available at https://github.com/liliancaesar/Publication_scripts/tree/main/2020_Longitudinal_survey/Statistical_analyses. Data were tested for their fit to normality and variance homogeneity using Shapiro’s and Bartlett’s tests (*P* < 0.05), respectively. When applicable, package MASS [26] was used for Box-Cox transformation of non-normal and non-homogeneous data. Temporal differences in forager weight were assessed by one-way ANOVA using “month” as factor. Monthly differences of daily variations in colony temperature and humidity (Δt and Δh) were tested with Kruskal-Wallis. To test whether unhealthy colonies were on average cooler and/or more humid, the March daily lowest temperatures and highest humidities were analysed by Mann-Whitney. The package Laercio [27] was used for mean comparisons with Tukey’s test, and PMCMRplus [28] for Nemenyi’s test. Forager CNRQ differences for *CYP450, PO and ApoLp* were evaluated by one-way ANOVA using “month” and “colony” as factors separately. Taking into account that the syndrome occurs in March, and that colonies have been monitored from January until April, a two-way ANOVA using either “colony”, “MD colonies”, “health status” or “intensity” combined with “month” as factors was also performed, enabling us to identify gene expression effects in specific periods during the course of our survey. Pearson’s correlation coefficients among all traits were calculated with package Hmisc [29].

## Results

### (a) Differentially expressed genes in diseased colonies

Transcriptome sequencing of D1, D2 and H foragers yielded 99,766,936, 102,034,731 and 135,982,124 single-end high quality-trimmed reads, respectively. From all reads, 87-99% mapped against the *M. quadrifasciata* genome. A total of 558 DEGs were found comparing bees from healthy and both diseased colonies, with 493 downregulated in foragers from diseased colonies (figure 1B; electronic supplementary material, table S2). Membrane components (GO:0016020 and GO:0016020) are significantly enriched in DEGs (figure 1C; electronic supplementary material, table S3), from which the vast majority is downregulated, representing major deficits in bees affected by the syndrome. To our knowledge, most of the highly differentially expressed genes were never directly implicated in bee health. Therefore, three DEGs homologous to genes commonly found differentially expressed in unhealthy bees from other species were chosen to quantify temporal variations in gene expression in foragers under semi-controlled conditions using RT-qPCR, *i.e.*, *CYP450*, *PO* and *ApoLp* (electronic supplementary material, table S4). *CYP450* and *ApoLp* are both downregulated in our transcriptomes from diseased colonies, but curiously *PO* showed inconsistent patterns of differential expression in D1 and to D2 relative to H.

### (b) Outcomes of the survey

Among the six colonies monitored monthly in our study, four manifested signs of the syndrome in March 2019. Colonies BP2 and BP3 manifested the strongest signs of disease, with high mortality of workers, and some bees showing tremors or paralysis. Their respective daughter colonies PA2 and PA3 showed less intense signs and were characterized as mildly diseased. We rule out the possibility that bees died due to lethal doses of agrochemicals, since residue analyses did not indicate contamination by agrochemical compounds in bees from diseased colonies (data not shown). MD colony pair BP1 and PA1 did not manifest health alterations.

### (c) Temporal variation in gene expression

There was a significant temporal variation in the transcription of *CYP450* (*P* = 0.009), with peaks in March in both foragers from healthy and diseased colonies, and in January only in healthy ones (figure 1D). Monthly variations in *CYP450, PO* and *ApoLp* gene expression showed similar interactions with the colony health status (*P =* 0.0103, 0.0152 and < 0.0005, respectively), and foragers from colonies that remained healthy during the outbreak period showed the highest expression of these three genes in January (figure 1D).

### (d) Changes in worker bee nutrition and colony microclimate

We found a marked reduction in forager weight from January until March (*P* < 0.0005; figure 2A), with the lowest average weight reached during the syndrome outbreak (March). From January to February we observed a sudden change in the pollen stored by worker bees, *i.e.*, its composition shifted from Myrtaceae (as *Eucalyptus* sp.) to Fabaceae (mainly *Mimosa bimucronata;* figure 2B). Bee weight reduction was accompanied by a reduction in the internal temperature of colonies (*r =* 0.63, *P =* 0.0088), and increase in humidity (*r =* −0.58, *P =* 0.0179; electronic supplementary material, figure S1). This pattern was more pronounced in colonies that became diseased during the outbreak, *i.e.*, MD colonies 2 (BP2 and PA2; *P* < 0.0005). Furthermore, temperature was lowest (*P* < 0.0005) and humidity was highest (*P* < 0.0005) inside diseased colonies when the syndrome symptoms were first observed in March (figure 2C). The highest differences in daily temperature and humidity within colonies occurred from December to March (*P* < 0.0005; figure 2D).

**Figure 2:**
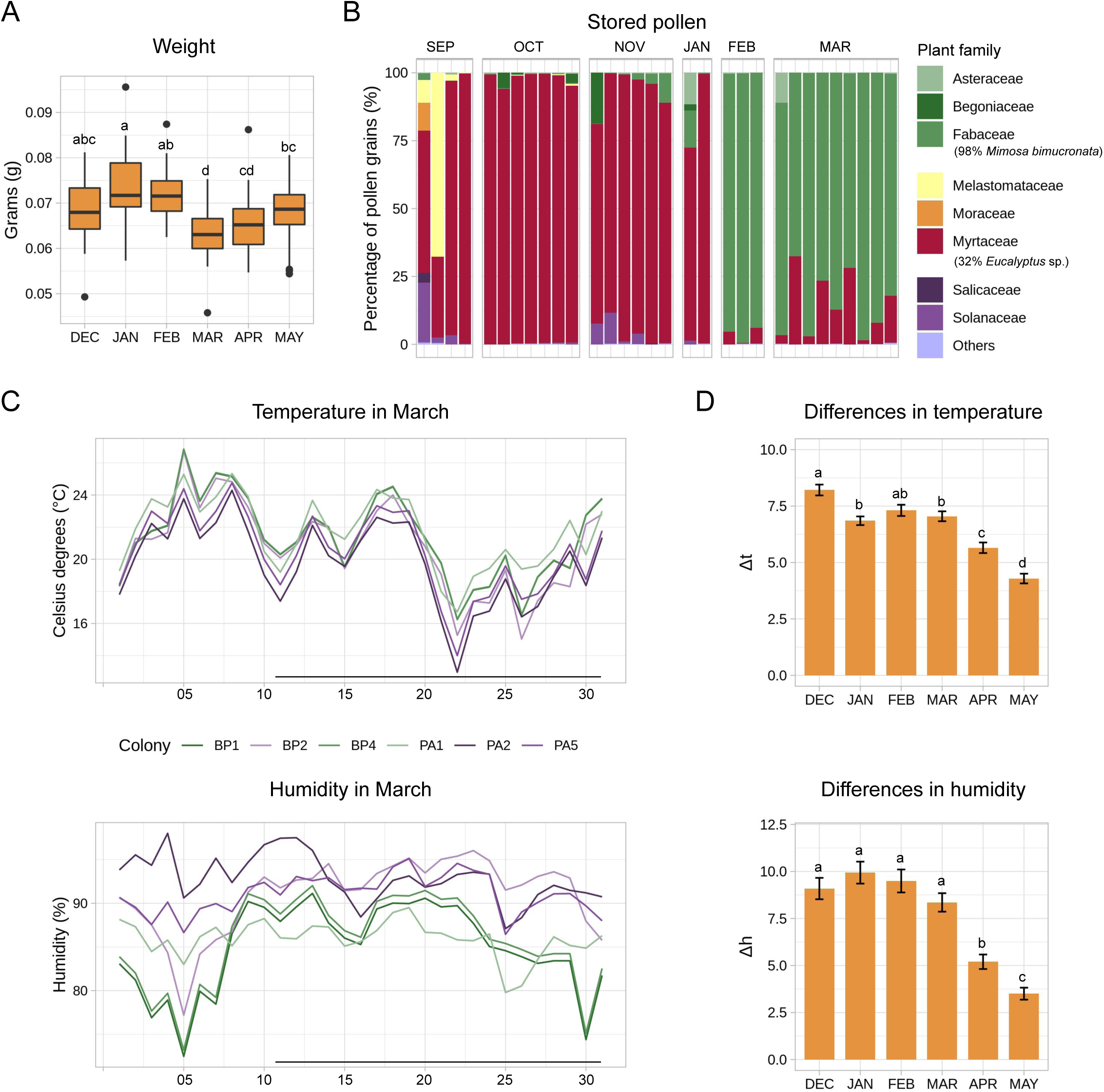
(A) Boxplot showing the monthly variation in average forager weight. (B) Barplot showing the monthly variation in stored pollen, revealing a switch between January and February. (C) Line plot of the minimum daily temperature and maximum daily humidity in March, in which the colors identify colonies that remained healthy (shades of green) or became diseased (shades of purple) during the outbreak period (horizontal bars). (D) Barplot showing the average daily difference in maximum *vs*. minimum colony temperature (Δt) and humidity (Δh). Vertical bars indicate the standard error; lowercase letters represent significant differences according to (A) Tukey’s test or (D) Nemenyi’s test (*P* < 0.05).

## Discussion

Our study revealed that both genetic and environmental factors influence the annual *M. quadrifasciata* syndrome. Mother-daughter colonies, exhibiting an average genetic relatedness of 0.375 [30], showed similar health status during the outbreak, despite having been kept in different localities. We suspect that the routine practice of colony division, associated with the concomitant loss of wild nests, may have reduced the genetic diversity of *M. quadrifasciata* populations. Reduced genetic diversity may have affected stingless bees to properly respond to environmental stresses in the form of sublethal effects such as pesticides, pathogens, and limited food resources, predisposing them to disease. We found that symptoms of worker bees were stronger in two affected colonies located nearby an agricultural setting. Thus, two lines of evidence allow us to identify some of the mechanisms involved in a higher susceptibility to disease, and eventually colony failure, possibly through the impairment of forager responses to interacting stressors.

Firstly, our exploratory transcriptomic analyses showed that components of the cellular membrane, representing the layer that directly communicates with the environment, are enriched with DEGs that are downregulated in foragers from colonies affected by disease. Furthermore, enrichment analysis also indicated a slight over representation of lipid metabolic processes and oxidoreductase molecular functions, which were previously shown to respond to pesticides in honey bees [7]. Secondly, foragers from our surveyed colonies affected by the syndrome in March expressed significantly less *CYP450, ApoLp and PO* in January. Bees rely on CYP450s for xenobiotic detoxification, which determines their sensitivity to agrochemicals [31]. ApoLp is a major component of the honey bee royal jelly [32] and is responsible for lipid transport, being mostly involved in innate immunity [33]. PO is an enzyme responsible for activating melanogenesis, an important defense mechanism of insects, and considered as an indicator of health condition strongly influenced by diet [34]. Thus, it is not surprising that the expression patterns of these three genes (specially *ApoLp* and *PO*) are so remarkably similar in our survey, suggesting the combined involvement of three physiological processes in the *M. quadrifasciata* syndrome, *i.e.*, xenobiotic detoxification, immunity and nutrition.

Agrochemicals are known to reduce ApoLp levels in honey bees [35], and increase their demand for food [36]. Pollen stress in turn may lead to developmental and behavioural impairment [6]. Moreover, January is characterized by a high density of pollinators in general, including *A. mellifera*, creating opportunities for getting in contact with a larger diversity of pathogens. In our study, pronounced forager weight loss occurred between January and March, as well as a shift in the pollen stored by worker bees from Myrtaceae to mostly *Mimosa bimucronata* between January and February. Such a shift may result from the opportunity to forage *Mimosa*, that starts to bloom in February, and from competitive exclusion, since *M. quadrifasciata* apparently competes with *A. mellifera* for Myrtaceae flowers [37]. Interestingly, two months after *M. quadrifasciata* initiated storing *M. bimucronata*, a gradual recovery of forager weight was noticed. Colony recovery after the outbreak period is also suggested by a better performance of worker bees in controlling their nest microenvironment, as observed by the reduced differences in colony daily temperature and humidity in April and May.

Unfortunately, we cannot rule out that the differences we observed in forager gene expression are due to age differences. *Melipona* spp. development takes around 40 days from egg to adult [38], thus *M. quadrifasciata* foragers sampled during the outbreak in March were immatures in January. However, age polyethism in bees is known to be a labile feature, since forager differentiation from nurses might be accelerated if the colony weakens [39]. Conducting invasive studies with stingless bees is not straightforward. Their colonies are much smaller than honey bees, semi-wild, and present a number of significant challenges for conducting experiments avoiding the side effects of excessive manipulation. Although four of the six colonies surveyed in our study manifested some degree of the syndrome symptoms, none of them collapsed during the outbreak period. We think that the annual collapses reported for *M. quadrifasciata* colonies in Southern Brazil could result from positive density dependence influenced by the combination of diverse sublethal factors [4,40]. The complexity of causes behind worldwide colony collapses demand efforts to sustain pollination services. Based on our findings, actions such as limiting the use of agrochemicals in the vicinity of managed colonies and providing abundant natural polyfloral resources through the conservation of native forests could help prevent the annual *M. quadrifasciata* syndrome. Furthermore, it warrants a better understanding of the impact of management practices on stingless bee genetic diversity and fitness.

## Acknowledgments

We thank the beekeeper Evald Gossler for making his meliponary available for the survey, and for collaborating throughout the whole survey. We also thank the beekeeper Gilson Maas for providing bee samples for our study.

