## Supplementary Figure S1 for "Longitudinal survey reveals delayed effects of forager gene expression on stingless bee colony health"

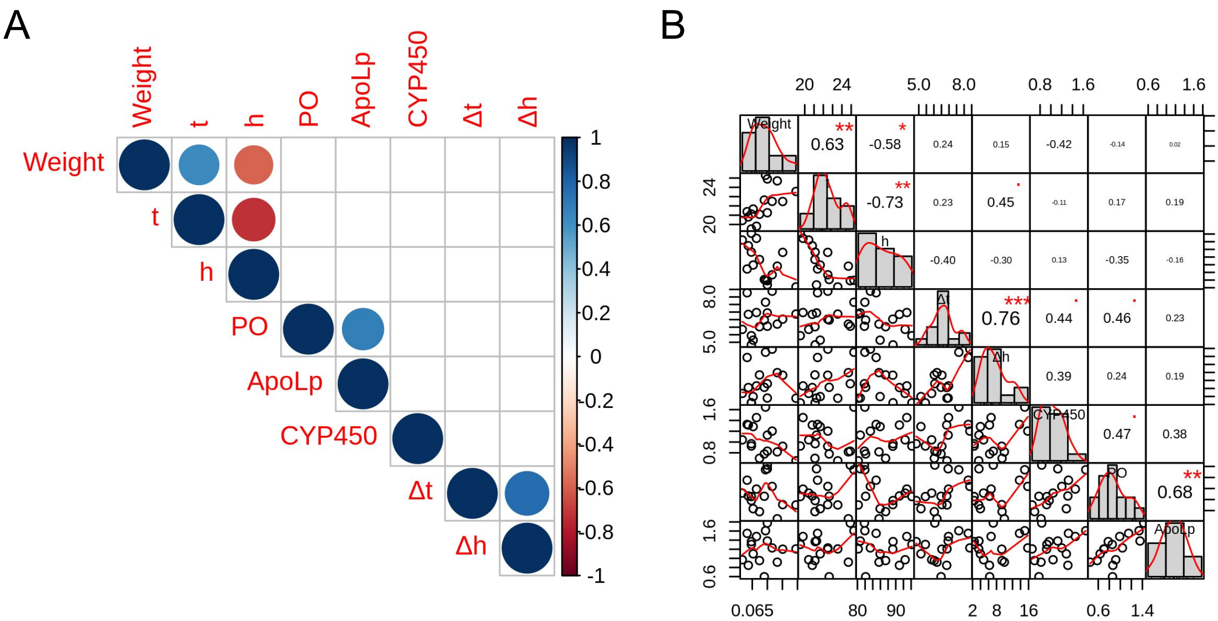


**Figure S1:** Pearson correlation matrix of all quantified traits at colony- and individual-level. (A) Significant positive and negative correlations are displayed in blue and red circles, respectively ($P$ < 0.05). Color intensity is proportional to the correlation coefficients ($r$), highlighted in the matrix on the right. (B) In addition to the correlation coefficient ($P$ codes '***' 0.001, '**' 0.01, '*' 0.05, '.' 0.1, ' ' 1), the matrix presents histograms with kernel density estimation, and scatter plots with fitted line.
