## Supplementary Text S1 for "Longitudinal survey reveals delayed effects of forager gene expression on stingless bee colony health"

**Electronic Supplementary Material, text S1**

**Detailed materials and methods**

**(a) Samples for transcriptome sequencing**

Two colonies were sampled during the outbreak of March 2018 in meliponaries from Bom Princípio (BP; 29°31′2.30′′S/51°17′29.00′′W) and Estância Velha (EV; 29°38'50.316''S/51°10'23.592''W). Both colonies were manifesting the syndrome signs, such as adult bee mortality, or bees presenting tremors and paralysis. A third colony, that showed no such signs during the outbreak period, was sampled in April (after the outbreak) in BP. Three bees from each colony (labeled as D1, D2 and H, respectively) were collected and stored at -80ºC until RNA extraction for transcriptome sequencing. Total RNA of individual bees was extracted with TRIzol™ Reagent (Thermo Fisher Scientific, USA), following the manufacturer's recommendations. RNA yield was assessed with Qubit fluorometer (Invitrogen, USA) and the integrity was checked on a 1% agarose gel. For sequencing, aliquots of 2 µg RNA treated with TURBO DNase (Thermo Fisher Scientific, USA) from three foragers were pooled with respective colony samples.

**(b) Transcriptome analyses**

Trimmomatic v.0.36 was run with default parameters to remove low quality reads [[1]](https://www.zotero.org/google-docs/?62Kmqv). Gene expression (*GE*) was estimated with *depth* command from Samtools v.1.3.1 [[2]](https://www.zotero.org/google-docs/?XFqnsd) by mapping the trimmed reads onto *M. quadrifasciata* genome (GenBank: GCA_001276565.1) with GSNAP v. 2018-07-04 [[3]](https://www.zotero.org/google-docs/?8AehB0) and recovering the number of reads per gene. To normalize gene expression (*NGE*) the following formula was used: *NGE = ( 100,000,000 / nr of mapped reads ) * GE*. Next, the similarity in gene expression (*SGE*) between transcriptomes was estimated by comparing *NGE* from healthy and diseased bee colonies with the following formula: *SGE = ( H + D ) / ( 2 * max )*. Where *H* and *D* are the *NGE*s from healthy and each diseased colony, respectively, and *max* is the largest *NGE* value from *H* and *D* being compared. The result ranges between 0.5 and 1, and genes were regarded as differentially expressed (DEGs) when *SGE* was equal or lower than 0.7 (All scripts available at https://github.com/liliancaesar/Publication_scripts/tree/main/2020_Longitudinal_survey/Transcriptome_analyses). Hypothetical genes among DEGs were re-annotated with BLASTp against the *nr* database (cutoff e-value 1e-5; electronic supplementary material, table S2) [[4]](https://www.zotero.org/google-docs/?i73zIy). To find functions over-represented in DEGs, a functional enrichment analysis was performed using the online version of g:Profiler with default parameters (g:SCS threshold of 0.05) and gene ontology annotation (GO terms) as data source [[5]](https://www.zotero.org/google-docs/?zPtsZj). DEGs with known roles for bee health were selected for relative quantification with RT-qPCR (see below). Protein annotations of selected DEGs were cross-checked with the online version of eggNOG-mapper v2 [[6]](https://www.zotero.org/google-docs/?eT20Uz), BLASTp analysis against *A. mellifera* proteins (taxid = 7460) and against the complete *nr* database of NCBI (cutoff e-value 1e-5; electronic supplementary material, table S4) [[4]](https://www.zotero.org/google-docs/?PxqdIF). Proteins were also compared to the Conserved Domains Database (CDD) using the Batch CD-Search tool (cutoff e-value 1e-3; electronic supplementary material, table S4) [[7]](https://www.zotero.org/google-docs/?ane2VB).

**(c) Monitoring bee colonies**

Surveyed colonies were monitored in two localities, *i.e.,* Bom Principio (BP; 29°31′2.30′′S/51°17′29.00′′W) and Porto Alegre (PA; 30°2'4.7292''S/ 51°13'3.5724''W). To control for genetic factors contributing to the syndrome, three colonies from BP (named BP1, BP2 and BP3) were divided in February 2018, resulting in three pairs of mother-daughter (MD) colonies. Daughter colonies (named PA1, PA2 and PA3) were translocated to PA after six months, when they became mature. All six colonies were monitored monthly at least from December 2018 (Summer) to May 2019 (Autumn).

**(d) Pollen resources used by stingless bees**

Aliquots of pollen stored by worker bees were collected with tweezers every month and stored in the laboratory at 4°C. When no stored pollen was found within our experimental hives, or storage was inaccessible for sampling, pollen was sampled from non-experimental colonies located nearby. Pollen was chemically processed by acetolysis [[8]](https://www.zotero.org/google-docs/?vTlhpB). Four slides were mounted for each sample with glycerin gelatine [[9]](https://www.zotero.org/google-docs/?woIJZC) and around 500 pollen grains were identified at the family, genus or species level using reference material from the pollen library of the Palynology Laboratory at Universidade Luterana do Sul do Brasil (Ulbra), ICN Herbarium at the Universidade Federal do Rio Grande do Sul (UFRGS), database of the Pollen Catalogs Network (RCPol; www.rcpol.org.br) and pollen descriptions of plants from Southern Brazil [[10–13]](https://www.zotero.org/google-docs/?Jcx4mE).

**(e) Relative quantification of gene expression**

With an entomological sucker, five foragers were collected monthly from each of the six colonies, between December 2018 and April 2019. They were weighted in digital precision scale, and transferred individually to vials containing 200 µL of RNAlater (Thermo Fisher Scientific, USA), followed by storage at -80°C for RNA extraction and gene expression analysis by RT-qPCR. Total RNA was extracted from the whole body and quantified as previously explained. An aliquot of 1 µg RNA from each forager was used as input for first strand cDNA synthesis with the High-Capacity Reverse Transcription kit (Thermo Fisher Scientific, USA). StepOnePlus™ Real-Time PCR System (Applied Biosystems) was used for the RT-qPCR assays. Amplifications were carried out in 25 μL reaction solutions containing 12.5 μL cDNA (diluted to 1:30), 0.2 X SYBR™ Green I Nucleic Acid Gel Stain (Thermo Fisher Scientific, USA), 0.25 U of Platinum Taq DNA polymerase (Invitrogen, USA), 1 X PCR buffer (Tris-HCl 200 mM, pH 8,4, KCl 500 mM), 3 mM MgCl2, 0.1 mM of each dNTP, 0.2 μM of each specific primer.

For template quantification we used the qBASE+ pipeline [[14]](https://www.zotero.org/google-docs/?x0vuiW), by first calculating the means and standard deviations of quantification cycle (Cq) values of technical replicates, and relativizing Cq values based on the gene specific amplification efficiency. Next, the sample specific normalization factors were calculated by taking the geometric mean of the relative quantities of the two reference genes (*act* and *rps5*). The normalized Cq values were finally rescaled in relation to the sample with the lowest relative quantity [[14]](https://www.zotero.org/google-docs/?nGJFi5), expressed in the form of calibrated normalized relative quantities (CNRQ), and used for statistical analyses.
